# A novel tomato inter-specific (*Solanum lycopersicum* var. *cerasiforme* and *S. pimpinellifolium*) MAGIC population facilitates trait association and candidate gene discovery in untapped exotic germplasm

**DOI:** 10.1101/2024.02.28.582481

**Authors:** Andrea Arrones, Oussama Antar, Leandro Pereira-Dias, Andrea Solana, Paola Ferrante, Giuseppe Aprea, Mariola Plazas, Jaime Prohens, María José Díez, Giovanni Giuliano, Pietro Gramazio, Santiago Vilanova

## Abstract

We developed a novel eight-way tomato multi-parental advanced generation inter-cross (MAGIC) population to improve the accessibility of the genetic resources of tomato relatives to geneticists and breeders. The inter-specific MAGIC population (ToMAGIC) was obtained by inter-crossing four accessions each of *Solanum lycopersicum* var. *cerasiforme* (SLC) and *S. pimpinellifolium* (SP), which respectively are the weedy relative and the ancestor of cultivated tomato. The eight exotic ToMAGIC founders were selected based on a representation of the genetic diversity and geographical distribution of the two taxa. The resulting MAGIC population comprises 354 lines which were genotyped using a new 12k tomato Single Primer Enrichment Technology (SPET) panel and yielded 6,488 high-quality SNPs. The genotyping data revealed a high degree of homozygosity (average 93.69%), an absence of genetic structure, and a balanced representation (11.62% to 14.16%) of the founder genomes. To evaluate the potential of the ToMAGIC population for tomato genetics and breeding, a proof-of-concept was conducted by phenotyping it for fruit size, plant pigmentation, leaf morphology, and earliness traits. Genome-wide association studies (GWAS) identified strong associations for the studied traits, pinpointing both previously identified and novel candidate genes near or within the linkage disequilibrium blocks. Domesticated alleles for fruit size were recessive and were found, at low frequencies, in wild/ancestral populations. Our findings demonstrate that the newly developed ToMAGIC population is a valuable resource for genetic research in tomato, offering significant potential for identifying new genes that govern key traits in tomato breeding. ToMAGIC lines displaying a pyramiding of traits of interest could have direct applicability for integration into breeding pipelines providing untapped variation for tomato breeding.

## Introduction

Tomato (*Solanum lycopersicum* L.) is the most economically important vegetable crop and a model plant species, with an extensive pool of genetic tools and resources. The tomato research community has access to a wealth of genetic information for wild species, landraces, and modern cultivars, including high-quality genome sequences (Rothan *et al.,* 2019). Several databases compiling genomic, genetic, transcriptomic, phenotypic, and taxonomic information are available (Fei *et al.,* 2006, 2010; Bombarely *et al.,* 2010; Suresh *et al.,* 2014; Kudo *et al.,* 2017). Over decades, several tomato bi-parental populations have also been released including introgression lines (ILs), recombinant inbred lines (RILs), advanced backcrosses (ABs), among others (e.g., Eshed and Zamir, 1995; Paran *et al.,* 1995; Tanksley and Nelson, 1996; Lippman *et al.,* 2007; Salinas *et al.,* 2013; Fulop *et al.,* 2016).

In the genomics era, new multi-parental populations have been developed dramatically increasing mapping resolution (Scott *et al.,* 2020). Multi-parent advanced generation inter-cross (MAGIC) populations are powerful next-generation pre-breeding resources with increased diversity and high recombination rates, suitable for QTL mapping and candidate gene identification (Mackay and Powell, 2007; Cavanagh *et al.,* 2008; Arrones *et al.,* 2020; Scott *et al.,* 2020). In tomato, only two MAGIC populations have previously been released. The first one was a MAGIC population developed by crossing four large-fruited *S. lycopersicum* accessions with four cherry-type accessions of *S. l.* var. *cerasiforme* (Pascual *et al.,* 2015). Final lines were used to study fruit weight distribution in the population in different environments, identifying QTLs that colocalized with already cloned genes. Subsequently, Campanelli *et al*. (2019) developed a MAGIC population that included seven cultivated accessions of tomato and one of the wild *S. cheesmaniae* as founders. The *S. cheesmaniae* accession was selected for its biotic and abiotic stress tolerance, yield and resiliency (Nesbitt and Tanskley, 2002).

The development of MAGIC populations using wild species as founders represents a promising way to combine the potential of these experimental populations for QTL/gene mapping together with the exploitation of the large phenotypic and genetic variation from the wild donor introgressions. Here, we present a novel eight-way inter-specific tomato MAGIC population (ToMAGIC) obtained by using *S. l.* var. *cerasiforme* (SLC) and *S. pimpinellifolium* (SP) accessions as founders, which respectively are the closest relative and the ancestor of cultivated tomato (Peralta *et al.,* 2008). Cultivated tomato suffered strong genetic bottlenecks during domestication and breeding processes, resulting in low genetic diversity of tomato landraces and heirlooms (Blanca *et al.,* 2015). Based on previous morphological characterization and resequencing data availability, the eight selected founders of the new ToMAGIC population represent a wide genetic and morphological variation, as well as differences in ecological adaptation (Blanca *et al.,* 2015; Gramazio *et al.,* 2020a). Founders are very diverse in terms of fruit, vegetative, and flowering traits but also their capacity of adaptation to different conditions, ranging from desert to tropical forest environments, and from sea level to over 1,500 m altitude. Therefore, one of the aims of this population is to recover Andean variability lost during the domestication process, by using a substantial proportion of the fully cross-compatible weedy and wild tomato diversity.

This ToMAGIC population may have a large potential to identify new genomic regions and candidate genes of interest in breeding, as well as to validate genes and QTLs already described in a genetic background other than that of cultivated tomato. In this way, another aim of this population is dissecting the control of different traits, including those involved in the early domestication of tomato (Frary and Doganlar, 2003). The introduction of exotic germplasm will be useful for shedding light on the genetics of agronomic and adaptation traits present in these materials, as well as for the selection of elite lines of interest for tomato breeding (Arrones *et al.,* 2020). In our work, the integration of high-throughput genotyping of the recombinant ToMAGIC population together with the phenotyping of specific traits across different plant parts has effectively demonstrated a proof-of-concept for the high-precision fine mapping of these traits. This approach has not only validated previously identified candidate genes for the traits studied in a SLC and SP genetic background, but also led to the discovery of new candidate genes, and the observation of additional phenotypic-causing variants, underscoring the great potential of the ToMAGIC population for tomato genetics and breeding.

## Materials and methods

### ToMAGIC founders

The inter-specific tomato MAGIC (ToMAGIC) population was developed through the inter-crossing of SLC and SP accessions. Founders consist of four weedy *S. l.* var. *cerasiforme*, i.e., BGV007931 (SLC1), LA2251 (SLC2), PI487625 (SLC3), and BGV006769 (SLC4), and four wild *S. pimpinellifolium,* i.e., BGV007145 (SP1), BGV006454 (SP2), BGV015382 (SP3), and BGV013720 (SP4). Their geographical origin, including geographical coordinates and altitude, and environmental parameters (Mean temperature, temperature range, precipitation, etc.) are known (Martínez-Cuenca *et al.,* 2020). With respect to the Heinz 1706 SL4.0 reference genome (Hosmani *et al.,* 2019), the total variants identified in SLC accessions ranged from 1.2 million in SLC2 to 1.9 million in SLC1, while in the SP accessions, they ranged from 3.1 million in SP4 to 4.8 million in SP3 (Gramazio *et al.,* 2020a). This set of variants was over 1,600-fold more abundant than the one used in the previous study of Blanca *et al*. (2015), where the eight founders were also genotyped.

### ToMAGIC population development

Although low heterozygosity levels were observed for founders in previous studies (Blanca *et al.,* 2015), before starting with the ToMAGIC population cross-design, two generations of selfing of the founders were performed to ensure high homozygosity. To develop the ToMAGIC population, founder lines were inter-crossed by following a “funnel” approach including two extra generations of inter-crosses among the offspring of the double hybrid crosses. These extra steps were performed to increase recombination events among the genomes of the eight founders during the population development to achieve better mapping and QTL identification resolution (Arrones *et al.,* 2020). The first step in developing the MAGIC population consistent in crossing the SLC parents with the SP ones to produce interspecific F_1_ hybrids (SLC1 × SP2, SLC2 × SP1, SLC3 × SP4, and SLC4 × SP3). These F_1_ hybrids were subsequently inter-crossed in pairs (SLC1 × SP2 with SLC2 × SP1, and SLC3 × SP4 with SLC4 × SP3) directly (’) and reciprocally (’’) to obtain four genetically segregating double hybrids (DHY1’, DHY1’’, DHY2’, and DHY2’’). In this way, genomes from both species were mixed since the beginning of the development of the MAGIC population. Then, DHY1’ or DHY1’’ individuals were crossed with DHY2’ or DHY2’’ individuals obtaining a set of materials coming from the first inter-cross generation (IC1), which were an admixture of the genomes of the eight founders. DHYs were crossed by following a chain pollination scheme, where each individual was used as a female and male parent of different crosses (Díez *et al.,* 2002; Mangino *et al.,* 2022). In the same way, individuals from the second inter-cross (IC2) generation were also inter-crossed following a chain pollination scheme. This step was repeated to obtain the individuals from the third inter-cross generation (IC3). Finally, progenies of the IC3 were selfed for five generations by single seed descent (SSD) to obtain the ToMAGIC recombinant inbred lines. To accelerate the obtention of the SSD generations, selfings were stimulated by mechanical vibration and pruning was done manually, regulating vegetative growth and flowering. A set of 354 ToMAGIC lines were used in this study for phenotyping and genotyping.

Seeds from the 354 ToMAGIC lines were germinated in seedling trays with Humin-substrat N3 substrate (Klasmann-Deilmann, Germany) in a climatic chamber under a photoperiod and temperature regime of 16 h light (25 °C) and 8 h dark (18 °C). Plantlets were subsequently transplanted to individual thermoformed pots (1.3 l capacity) for acclimatisation and grown in a pollinator-free glasshouse of the Universitat Politècnica de València (UPV, Valencia, Spain). Plants were fertirrigated using a drip irrigation system and trained with vertical strings. Phytosanitary treatments against whiteflies and *Tuta absoluta* were performed when necessary.

### High-throughput genotyping

Young leaf tissue was sampled from the 354 ToMAGIC lines. Genomic DNA was extracted using the SILEX extraction method (Vilanova *et al.,* 2020). DNA quality and integrity were checked by agarose electrophoresis and NanoDrop ratios (260/280 and 260/230), while its concentration was estimated using a fluorescent DNA intercalating agent (e.g., Quant-iT PicoGreen dsDNA Assay Kit, Thermo Fisher Cat. No. P7589) and a microplate reader (Thermo Fisher Scientific)). Samples were sent to IGATech company (Udine, Italy) for library preparation and sequencing (150 paired-end) for a high-throughput genotyping using a newly developed 12k probes tomato Single Primer Enrichment Technology (SPET) panel, which is considerably improved over the original 5k probes tomato set (Barchi *et al*., 2019). The new SPET panel comprises 12,000 probes and was developed by selecting the most informative and reliable polymorphisms (of which ∼11,500 within 100 nt of a gene and ∼500 in intergenic regions) (Aprea *et al*., in preparation).

Cleaning of raw reads was performed using Fastp (Chen, 2023). Clean reads were mapped onto the tomato reference genome Heinz 1706 SL4.0 (Hosmani *et al.,* 2019) using BWA-MEM (Li, 2013) with default parameters; finally, GATK was used for variant calling (DePristo *et al.,* 2011), following the best practices recommended by the Broad Institute. The SNPs identified by the tomato SPET panel were first filtered by coverage ≥ 10 and quality GQ ≥ 20 using bcftools (https://doi.org/10.1093/gigascience/giab008), and then filtered using the TASSEL software (ver. 5.0, Bradbury *et al.,* 2007) to retain the most reliable ones (minor allele frequency > 0.01, missing data < 0.1, and maximum marker heterozygosity < 0.7). In addition, a linkage disequilibrium (LD) k-nearest neighbour genotype imputation method (LD KNNi) was performed to fill the missing calls or genotyping gaps (Troyanskaya *et al.,* 2001). Final marker density along chromosomes was represented using the R package chromPlot (Oróstica and Verdugo, 2016).

### Population diversity analysis

A principal component analysis (PCA) was performed to assess the population structure of the MAGIC population. PCA scores were generated in TASSEL software (ver. 5.0, Bradbury *et al.,* 2007). For graphically plotting the final PCA results the R package ggplot2 was used (Wickham, 2016). A heat map of the kinship matrix to identify possible relationships between lines was generated with GAPIT software (v.3, Wang and Zhang, 2021). A dendrogram of the MAGIC population was generated using the neighbor-joining method (Saitou and Nei, 1987) and the graphical representation was displayed and edited using the iTOL v.4 software (Letunic and Bork, 2019) to evaluate the genetic similarities among ToMAGIC lines and founders. Parental contribution to the ToMAGIC lines and haplotype blocks was estimated by using the R package HaploBlocker (Pook *et al*., 2019).

### ToMAGIC phenotyping

A proof-of-concept for testing the potential of the MAGIC population for GWAS analysis and detection of genomic regions associated with different types of traits was performed by phenotyping the eight parents and the 354 ToMAGIC lines for a set of traits from different plant organs. The traits evaluated included two related to fruit size (fruit locule number and fruit weight), one to plant pigmentation (plant anthocyanin), two to leaf morphology (lobing/serration and leaf complexity), and one to earliness (number of leaves below the first inflorescence). Tomato fruits evaluated for fruit weight and cut transversally for locule number counting. Presence of plant anthocyanin was observed in vegetative plant parts (stem, branches, leaf veins or leaf area) and scored in a range from 0 (slight presence, mainly on the stem) to 4 (strong presence in all plant parts). Leaf lobing/serration was scored in a range from 1 (lack of lobing/serration) to 7 (very serrated leaf). Leaf complexity was screened using a binary classification for pinnate (0) and bipinnate (1) compound leaves. The number of leaves below the first inflorescence was recorded by counting the leaves of the primary shoot when the first flower bud was visible. Pearson pair-wise coefficient of correlation (r) among traits was calculated, and their significance was assessed using a Bonferroni correction at the p<0.05 probability level (Hochberg, 1988) using R packages psych (Revelle, 2007) and corrplot (Wei and Simko, 2017).

### Genome-Wide Association Study (GWAS)

Using the genotypic and phenotypic data collected from the ToMAGIC lines, GWAS analyses were performed for the selected traits using the GAPIT software (v.3, Wang and Zhang, 2021). General linear model (GLM), mixed linear model (MLM), and BLINK analyses were conducted for the association study (Price *et al.,* 2006; Yu *et al.,* 2006; Huang *et al.,* 2019). Comparison of models was displayed in roundness Manhattan plots and QQ plots. The multiple testing was corrected with the Bonferroni and the false discovery rate (FDR) methods (Holm, 1979; Benjamini and Hochberg, 1995) with a significance level of 0.05 (Thissen *et al.,* 2002). SNPs with a limit of detection (LOD) score (calculated as -log10[p-value]) exceeding these specified thresholds or cutoff values in the three GWAS models were considered significantly associated with the traits under evaluation. Associations were considered significant if the same SNP exceeded the cut-off thresholds in at least two of the implemented models, indicating robustness. The top significant SNPs and their neighboring SNPs were used to calculate the correlation coefficient (r^2^). SNPs with default r^2^ values greater than 0.5 were considered for haplotype block estimation. The R package geneHapR was used for haplotype statistics (Zhang *et al.,* 2023a). The genes underlying the haplotype blocks were retrieved from the Heinz 1706 SL4.0 tomato reference genome (Hosmani *et al.,* 2019). Genes were considered as potential candidates in controlling the assessed traits according to SnpEff software v 4.2 prediction (Cingolani *et al.,* 2012) of the eight MAGIC founders (Gramazio *et al.,* 2019). The Integrative Genomics Viewer (IGV) tool was used for the visual exploration of founder genome sequences to validate SnpEff results (Robinson *et al.,* 2023). A conservative domain analysis was performed using the NCBI conserved domain server (https://www.ncbi.nlm.nih.gov/Structure/cdd/wrpsb.cgi) to assess the predicted variants at the protein level. The BLASTp (e-value cut-off of 1e^-10^) alignment tool and EnsemblPlants browser were used to compare the homology of protein sequences encoded by genes belonging to the same gene family. Haplotype and phenotype boxplots and density plots were generated with the R package ggplot2 (Wickham, 2016). To assess the significance of differences among different haplotypes pairwise *t*-tests were performed.

## Results

### MAGIC population construction

In the first stage of MAGIC population development, SLC and SP accessions of different origins (Figure 1A) were inter-crossed pairwise (Figure 1B). These materials are native to different geographic regions of South and Central America, mainly from Ecuador and Northern Peru, and provide a representation of the Andean variability lost during the domestication process in Mesoamerica (Figure 1A). They were selected since they are considered genetic diversity reservoirs barely exploited in tomato breeding (Gramazio *et al.,* 2020a). They include a wide molecular variability and phenotypic diversity in plant and inflorescence architecture, leaf, flower, and fruit traits, together with resistance or tolerance - in some of the founders - to biotic and abiotic stresses (Blanca *et al.,* 2015), including water and salt stress adaptation (Martínez-Cuenca *et al.,* 2020). The eight founders have previously been characterized morphoagronomically and their genomes have been resequenced (Blanca *et al.,* 2015; Gramazio *et al.,* 2020a).

**Figure 1.**
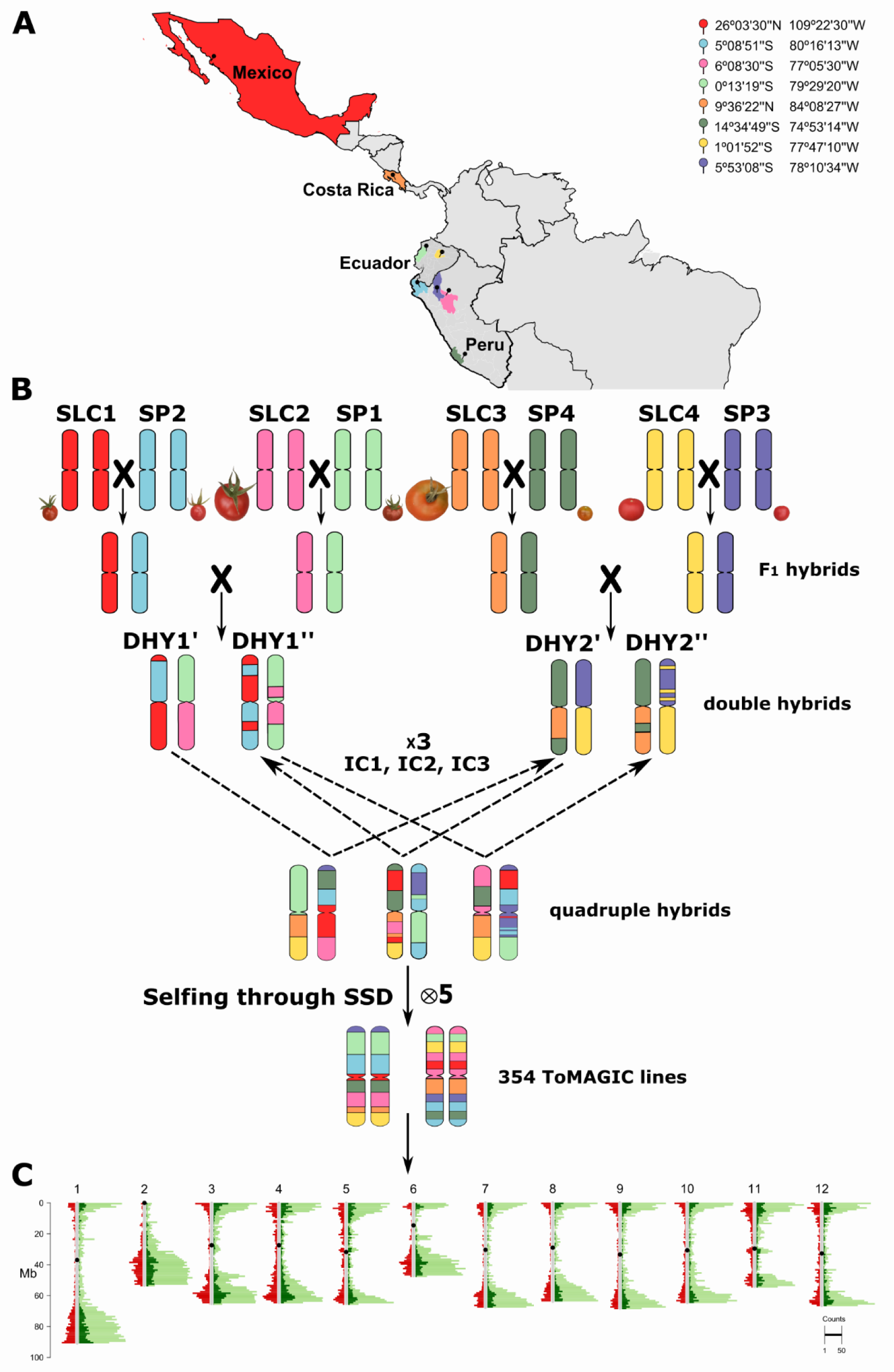
(A) Origin of the different SLC and SP founders selected for the ToMAGIC population development represented with the different colours code. (B) The funnel breeding design to develop the 354 ToMAGIC lines. The eight founders with a different colour to represent their genomic background, are represented at a scale based on the real fruit size. (C) Distribution of the 6,488 filtered markers (in red), the Heinz 1706 SL4.0 annotated genes (in light green), and the genes covered by the filtered markers (in dark green) across the 12 tomato chromosomes.

These weedy (SLC) and wild (SP) tomato species are cross-compatible (Gramazio *et al.,* 2020a), and thus the manual inter-cross was successfully performed. As a result of the inter-cross of the eight founders, the F_1_ hybrids, and the DHY hybrids, 112 IC1 individuals were obtained. The subsequent inter-crossing following a chain pollination scheme resulted in the obtention of 232 IC2 and 481 IC3 individuals. The latter individuals were self-pollinated to produce 475 S1, 452 S2, 427 S3, 400 S4, and the final population of 354 S5 (ToMAGIC) lines (Figure 1B).

### Genotyping

A total of 4,268,587 SNPs were generated from the genotyping of the 354 ToMAGIC lines using a newly developed 12k probes tomato SPET panel (Aprea *et al*., in preparation). After filtering, 6,488 markers were retained for the subsequent GWAS analysis. A higher marker density was observed in gene-rich regions located in distal chromosomal regions (Figure 1C). The distribution of SNPs among the different tomato chromosomes was fairly uniform, with an average marker density of 8.51 per Mb (Table 1). There was a marker average interval of 0.13 Mb with the broader marker intervals being around the pericentromeric regions. The filtered markers cover 16.91% of the total annotated genes. The residual heterozygosity of the ToMAGIC lines was on average 6.31%.

**Table 1.** Chromosome-wide distribution of the SNP positions used for the genome-wide association study (GWAS) in the tomato MAGIC population.

| Chromosome | Markers | % Markers | Chromosome length (Mb) | Marker density (markers/Mb) | Marker interval (Mb) |  | Genes | Covered genes |
| --- | --- | --- | --- | --- | --- | --- | --- | --- |
|  |  |  |  |  | Max. | Average |  |  |
| 1 | 646 | 10.78 | 90.86 | 7.11 | 3.60 | 0.14 | 4,133 | 619 |
| 2 | 568 | 7.25 | 53.47 | 10.62 | 3.77 | 0.09 | 3,379 | 518 |
| 3 | 624 | 8.30 | 65.30 | 9.56 | 2.52 | 0.10 | 3,324 | 578 |
| 4 | 815 | 8.75 | 64.46 | 12.64 | 1.32 | 0.08 | 2,819 | 689 |
| 5 | 656 | 8.44 | 65.27 | 10.05 | 1.57 | 0.10 | 2,382 | 496 |
| 6 | 408 | 7.84 | 47.26 | 8.63 | 3.11 | 0.11 | 2,769 | 392 |
| 7 | 425 | 8.07 | 67.88 | 6.26 | 2.97 | 0.16 | 2,517 | 379 |
| 8 | 346 | 8.14 | 64.00 | 5.41 | 3.35 | 0.19 | 2,428 | 315 |
| 9 | 436 | 8.02 | 68.51 | 6.36 | 4.26 | 0.16 | 2,521 | 403 |
| 10 | 406 | 7.46 | 64.79 | 6.27 | 2.58 | 0.16 | 2,52 | 342 |
| 11 | 552 | 7.51 | 54.38 | 10.15 | 1.69 | 0.10 | 2,326 | 446 |
| 12 | 606 | 9.43 | 66.69 | 9.09 | 1.97 | 0.11 | 2,444 | 498 |
| Total | 6,488 | 100 | 772.87 |  |  |  | 33,562 | 5,675 |
| Average |  | 8.33 |  | 8.51 |  | 0.13 |  |  |

### Population structure

A lack of genetic structure in ToMAGIC population was supported by the Principal Component Analysis (PCA), in which no differentiated groups were observed (Figure 2A). The first two PCs accounted only for 3.40% of the genetic variance, the first ten PCs 9.93%, and it required 41 PCs to explain 20% of the genetic variation, underscoring the weak population structure of the population. In addition, kinship coefficients between pairs of ToMAGIC lines varied from 0 to 1.32 (on a scale of 0 to 2), with 98.35% of the pairs with kinship values <0.5 (Figure 2B). These results revealed a low genetic relatedness among ToMAGIC lines.

**Figure 2.**
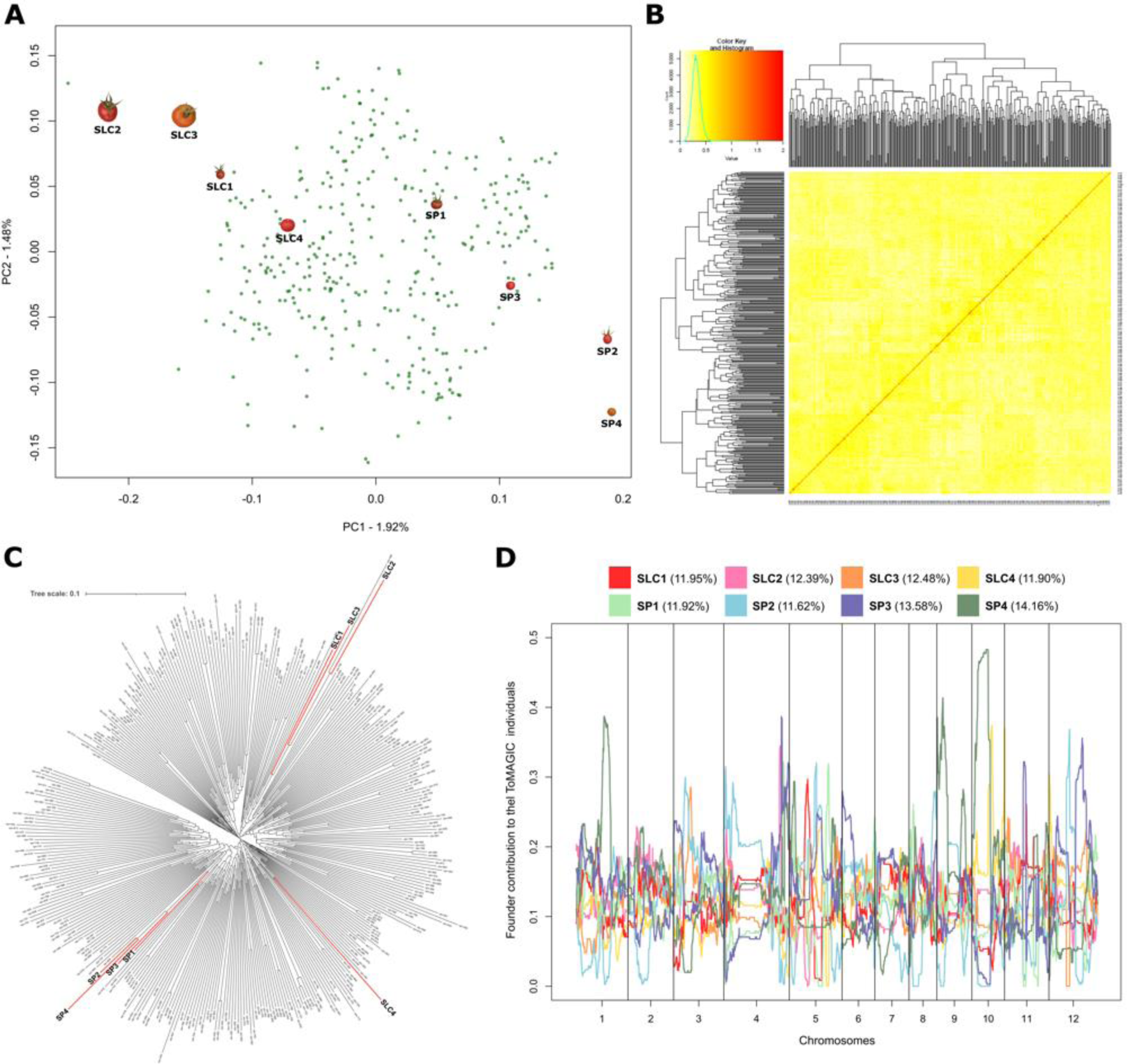
Population structure of the inter-specific ToMAGIC population. (A) Principal component analysis (PCA) plot of the first two PCs. (B) Heatmap plot of genetic relationship based on the kinship matrix. (C) Dendrogram indicating founders’ locations with coloured red branches. (D) Genome-wide founder haplotype blocks assignment across the 12 tomato chromosomes (x-axis) as the average percentage of founders’ contribution to the ToMAGIC lines (y-axis) with a different colour associated with each founder.

SLC founders were grouped close together, with negative values of the PC1, while SP founders had positive values for the PC1 (Figure 2A). A similar grouping was observed in the dendrogram of the MAGIC population and founders (Figure 2C). SLC2 and SLC3 are the closest accessions to cultivated tomato and plot in the first PCA quadrant with low values for the PC1 and high for the PC2. SLC4 is the closest to SP founders in the PCA (Figure 2A) and is separated from the rest of SLC founders in the dendrogram (Figure 2C). The estimated average contribution of each founder to the overall population was around the theoretically expected value of 12.50%, with the range varying from 11.62% for SP2 to 14.16% for SP4. However, the reconstruction of genome mosaics for the 354 ToMAGIC lines, considering the eight founder haplotypes, revealed different haplotype block proportions at different chromosomal positions (Figure 2D).

### Phenotyping analysis

Phenotyping for locule number, fruit weight, plant anthocyanin pigmentation, leaf lobing/serration, leaf complexity, and number of leaves below the first inflorescence revealed a wide range of variation, including transgressive lines for some of the studied traits (Table 2, Figure S1A). For the locule number trait, the average for SP founders was 2 lobules, while the average for SLC was 2.75 and the range between 2 and 4. However, ToMAGIC lines with up to 5 and 6 locules were identified, although most of the lines only had 2 locules, resulting in an average value of 2.2. For the fruit weight, ToMAGIC lines showed an intermediate average (2.72 g) between the SP and SLC founders weight averages of 1.60 g and 4.97 g, respectively. However, the range of variation of the founders was greater (from 0.97 g to 11.59 g) than those of the ToMAGIC lines (0.44 to 7.01), and no lines were found with a higher weight than the heaviest founder (SLC3). For the plant anthocyanin pigmentation, the mean of SLC founders (0.50) was lower than that of the SP founders (1.25), mainly due to the high level of plant pigmentation of the SP4 founder. The range of variation was greater for the ToMAGIC lines (from 0 to 4) than for the founders (from 0 to 3). For the leaf lobing/serration, ToMAGIC lines showed an intermediate average (3.69 g) between the SP and SLC founders averages of 2.50 and 6, respectively. The ToMAGIC lines covered all the variation range found in the founders, from the lack of lobing/serration (1) to very serrated leaves (7). For the leaf complexity, ToMAGIC lines showed an intermediate average (0.26) between the SP (0) and SLC (0.50) founders. For the number of leaves below the first inflorescence, the SP founders had a slightly lower number (4.33) than SLC founders (6.66), while ToMAGIC lines had an average of 5.36 leaves. However, the range of variation was much larger for the ToMAGIC lines (from 4 to 10) than for the founders (from 4 to 7). Pearson pairwise correlations among the traits evaluation were conducted, and only a slight positive correlation (r = 0.3261; p=1.57e^-7^) between leaf lobing/serration and leaf complexity, was observed (Figure S1B).

**Table 2.** Means and range values for SLC and SP founders and ToMAGIC lines for the phenotypic traits evaluated.

| Trait | SLC |  | SP |  | ToMAGIC lines |  |
| --- | --- | --- | --- | --- | --- | --- |
|  | Average | Range | Average | Range | Average | Range |
| Locule number | 2.75 | 2 - 4 | 2 | 2 | 2.20 | 2 - 6 |
| Fruit weight | 4.97 | 1.61 - 11.59 | 1.60 | 0.97 - 2.89 | 2.72 | 0.44 - 7.01 |
| Plant anthocyanin | 0.50 | 0 - 1 | 1.25 | 0 - 3 | 0.94 | 0 - 4 |
| Leaf lobing and serration | 6 | 5 - 7 | 2.50 | 1 - 3 | 3.69 | 1 - 7 |
| Leaf complexity | 0.50 | 0 - 1 | 0 | 0 | 0.26 | 0 - 1 |
| Number of leaves below the first inflorescence | 6.66 | 6 - 7 | 4.33 | 4 - 5 | 5.36 | 4 - 10 |

### Fruit size

#### Locule number

The Manhattan plot for fruit locule number revealed one significant peak on chromosome 2 (Figure 3A, Table 3). For the GLM model, 25 SNPs were above the FDR threshold (LOD > 4.15), 20 of them over the Bonferroni threshold (LOD > 5.11) between 44.78 and 46.13 Mb. For the MLM model, 15 SNPs were above the FDR threshold, nine of them over the Bonferroni threshold between a reduced region of 44.82 and 46.02 Mb (Figure 3B). For the BLINK model, a single SNP was above the FDR and Bonferroni thresholds (LOD = 15.27) at 45.87 Mb position. This association peak accounted for 26.84% of the total phenotypic variance of the locule number trait.

**Figure 3.**
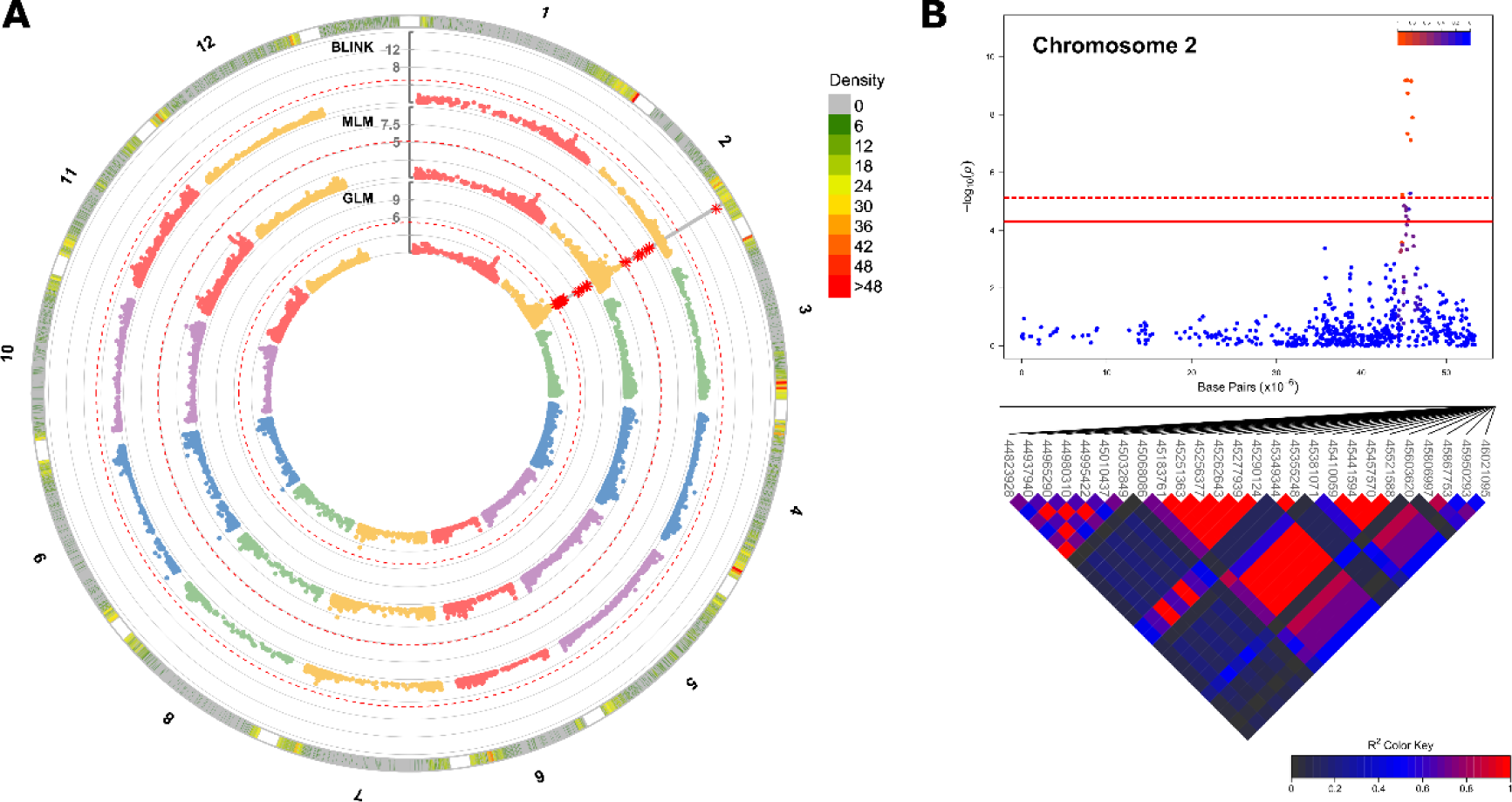
Genome-wide association results for the locule number trait. (A) Manhattan plots comparing GLM, MLM and BLINK models. The solid grey line indicates the common significant markers detected by two or more models. The red asterisks indicate the SNPs exceeding the Bonferroni threshold, represented as a dashed red line. (B) On the top, a chromosome-wise Manhattan plot with the top significant markers. Bonferroni and FDR thresholds are represented with red dashed and continuous lines, respectively. The colour from blue to red indicates r^2^ from 0 to 1. On the bottom, heat map of pairwise linkage disequilibrium (LD). SNP positions under the significant region are indicated in bp. The colour from black to red indicates r^2^ from 0 to 1.

**Table 3.**
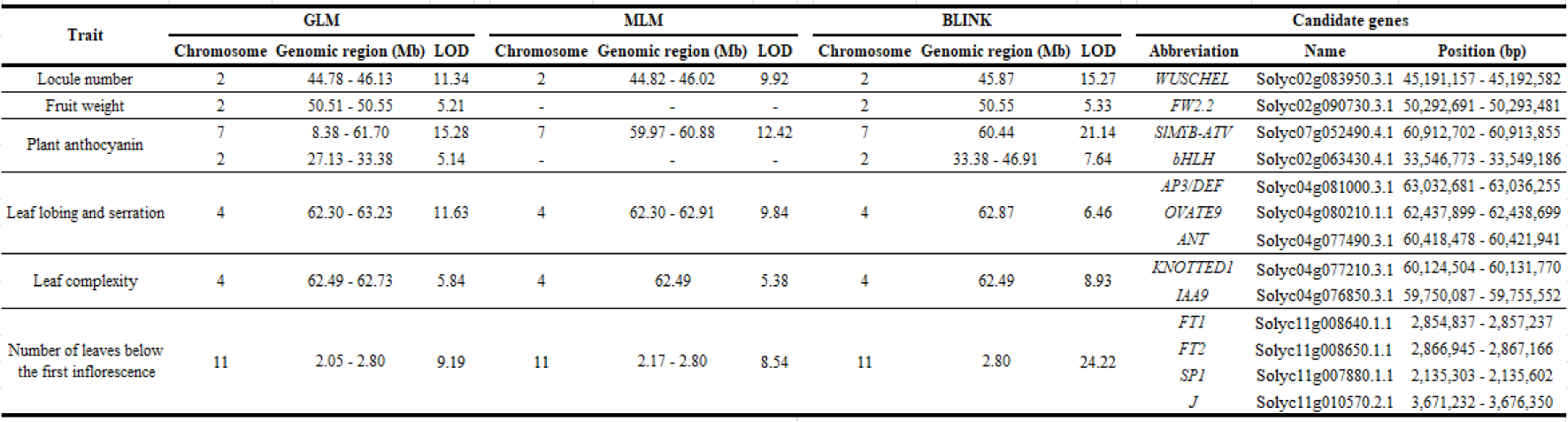
Association analysis results for GLM, MLM, and BLINK models and list of candidate genes for locule number, fruit weight, plant anthocyanin, leaf lobing/serration, leaf complexity, and number of leaves below the first inflorescence.

In the genomic candidate region on chromosome 2, the *WUSCHEL* gene (Solyc02g083950.3.1, 45,191,157-45,192,582 bp) was identified (Table 3). *WUSCHEL* gene controls stem cell fate in the apical meristem directly affecting locule number during tomato fruit development (Barrero *et al.,* 2006; Muños *et al.,* 2011). The two multi-locular founders of the ToMAGIC population, SLC2 and SLC3, showed two SNPs immediately downstream of the *WUSCHEL* gene that were previously described as directly associated with an increased locule number (Muños *et al.,* 2011). Specifically, a T/C transition at 45,189,386 bp and a A/G transition at 45,189,392 bp are considered as the responsible SNPs for the locule number trait (Figure S1A). These two SNPs were in almost complete linkage disequilibrium, and they are considered as a unique haplotype.

Haplotype analyses were performed to associate the candidate genomic regions with the phenotypic effects. For the locule number, a significant difference was observed between the haplotype of the SLC2 and SLC3 founders, which are the ones showing more than 2 locules, and the rest of the haplotypes of the ToMAGIC founders according to pairwise *t*-test for multiple comparisons (Figure 4A). When generating the density plot, higher values were also associated with the SLC2 (at 3 locules) and SLC3 (at 4 locules) founder haplotype.

**Figure 4.**
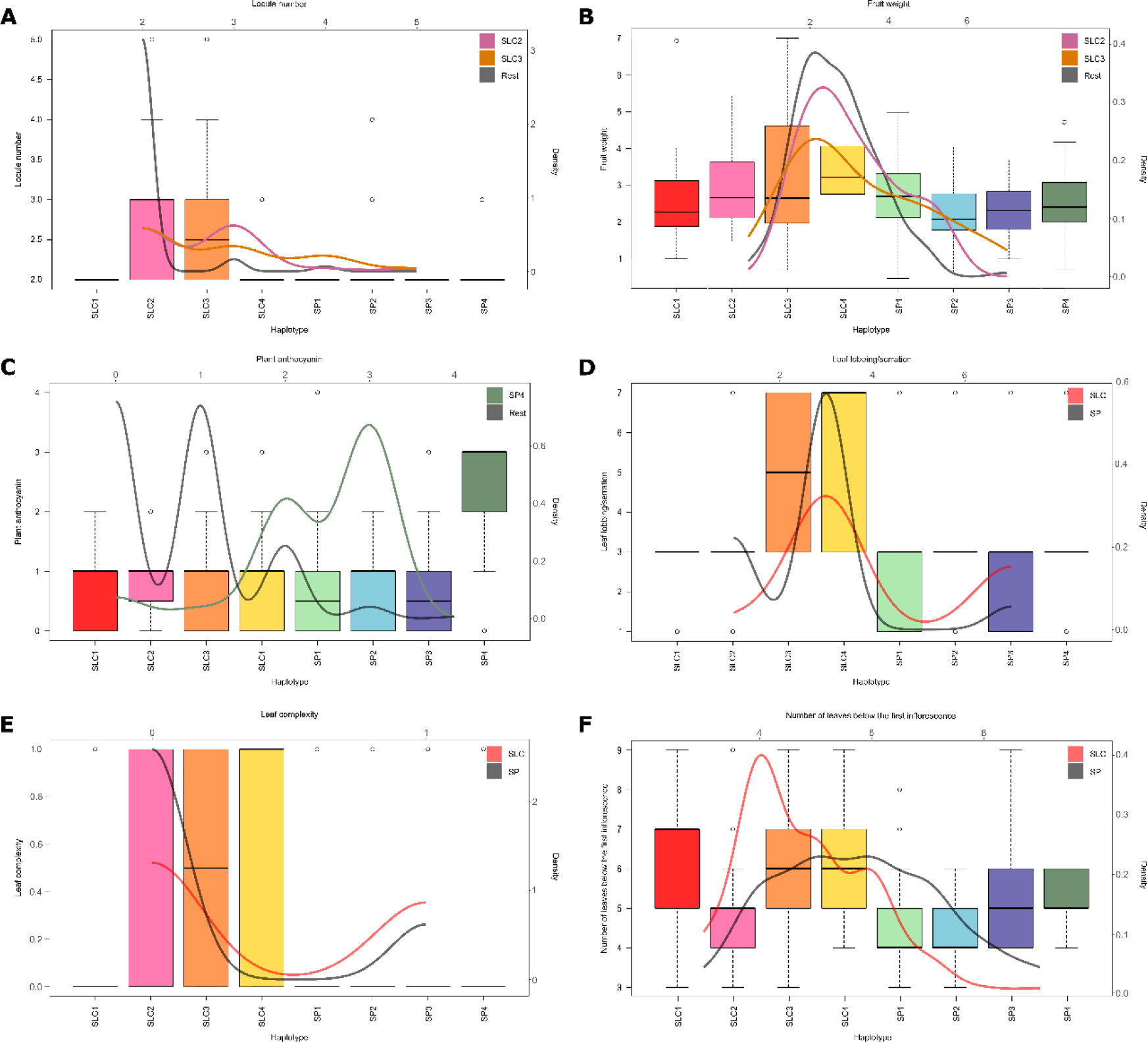
Haplotype analysis of the ToMAGIC lines for each of the MAGIC founders’ haplotype in combination with phenotypic data. Boxplot and density plot distribution in the candidate genomic regions for: (A) locule number; (B) fruit weight; (C) plant anthocyanin on chromosome 7; (D) leaf lobing/serration; (E) leaf complexity; and (F) number of leaves below the first inflorescence.

#### Fruit weight

The Manhattan plot for fruit weight also revealed one significant peak on chromosome 2, although only for GLM and BLINK models (Figure S2A, Table 3). For the GLM model, three peaks were above the Bonferroni threshold (LOD > 5.11) between

50.51 and 50.55 Mb (Figure S2B). For the BLINK model, a single SNP was above the Bonferroni threshold (LOD = 5.33) at 50.55 Mb position. This association peak explained 14.76% of the total phenotypic variance of the fruit weight trait.

Under the significant peak on chromosome 2, the well-known *FW2.2* gene (Solyc02g090730.3.1, 50,292,691-50,293,481 bp) was identified (Table 3). This gene is differentially expressed in floral development and controls carpel cell division (Frary *et al.,* 2000). The wild-type SNP was identified in all the ToMAGIC founders, except for founders SLC2 and SLC3, which have larger fruit weights (Blanca *et al.,* 2015). This SNP corresponds to a C/T change upstream of the 5’ region of *FW2.2* gene at 50,292,019 bp (Figure S1A).

In the haplotype analysis, pairwise *t*-test revealed a significant difference between SLC2 and SLC3 on one side and SP founders from the other (Figure 4B). When generating the density plot, most of the lines are around 2 to 3 g since light fruits predominate in the ToMAGIC population with an average weight of 2.72 g (Table 2). Lines with weights greater than 3 show mostly SLC2 and SLC3 haplotypes.

### Plant pigmentation

The Manhattan plot for plant anthocyanin revealed two significant peaks: one major peak on chromosome 7 and one minor but significant peak on chromosome 2 (Figure S3A, Table 3). For the GLM model, 31 SNPs were above the FDR threshold (LOD > 3.80) on chromosome 7, 21 of them over the Bonferroni threshold (LOD > 5.11) between 8.38 and 61.70 Mb. On chromosome 2, only four SNPs were above the FDR threshold, being two of them over the Bonferroni threshold between 27.13 and 33.38 Mb. For the MLM model, only one association peak was identified on chromosome 7 with eight SNPs over the FDR threshold, five of them over the Bonferroni threshold between a reduced region of 59.97 and 60.88 Mb (Figure S3B). For the BLINK model, a single SNP was above the FDR and Bonferroni thresholds (LOD = 21.14) on chromosome 7 at 60.44 Mb position. On chromosome 2, only two SNPs were above the FDR and Bonferroni thresholds at 33.38 and 46.91 Mb positions (LOD = 7.64 and 6.70, respectively). The association peak on chromosome 7 explained 15.14% of the total phenotypic variance of the plant anthocyanin trait, while the peak on chromosome 2 explained 4.68% of the phenotypic variance.

Under the major GWAS peak on chromosome 7, in the genomic region of 60,912,702-60,913,855 bp, a *MYB-like* transcription factor (*SlMYB-ATV,* Solyc07g052490.4.1) was identified (Table 3). The *SlMYB-ATV* (myeloblastosis-atroviolacea) gene has been described as a repressor of anthocyanin synthesis in vegetative tissues of tomato plants (Colanero *et al.,* 2018). However, we did not observe the previously described mutations in the gene sequence in our accessions. In contrast, a 9-bp in frame deletion at 60,912,903 bp position, deleting 3 amino acids in the transcriptional repressor MYB domain was identified in the SP4 founder, which is the unique founder showing anthocyanins in all plant parts (Figure S1A).

The same procedure was followed for the minor peak on chromosome 2. All the genes located near or within the LD block were assessed by SnpEff (Cingolani *et al.,* 2012) for all of the MAGIC founders. However, no potential candidate genes were identified, since no high-effect variants were predicted distinguishing between anthocyanin-containing and anthocyaninless founders.

In the haplotype analysis for chromosome 7, a significant difference was observed between the SP4 founder, which is the one showing increased levels of plant anthocyanins, and the rest of ToMAGIC founders according to pairwise *t*-test (Figure 4C). When generating the density plot, higher anthocyanin values were also associated with the SP4 founder haplotype.

### Leaf morphology

#### Leaf lobing/serration

The Manhattan plot for leaf lobing/serration revealed one significant peak on chromosome 4 (Figure S4A, Table 3). For the GLM model, 13 SNPs were above the FDR threshold (LOD > 4.81), ten of them over the Bonferroni threshold (LOD > 5.11) between 62.30 and 63.23 Mb. For the MLM model, ten SNPs were above the FDR threshold, nine of them over the Bonferroni threshold between a reduced region of 62.30 and 62.91 Mb (Figure S4B). For the BLINK model, a single SNP was above the FDR and Bonferroni thresholds (LOD = 6.46) at 62.87 Mb position. This association peak accounted for 53.84% of the total phenotypic variance of the leaf lobing/serration trait.

Different genes involved in the leaf shape were detected within the candidate genomic region on chromosome 4 identified in the GWAS for the leaf lobing/serration (Table 3). In order of proximity to the candidate region, we found the *APETALA3/DEFICIENS* or *AP3/DEF* gene (Solyc04g081000.3.1 between 63,032,681-63,036,255 bp), which has been described as a regulator of petal and sepal development (Quinet *et al.,* 2014), the ovate family protein 9 or *OVATE9* gene (Solyc04g080210.1.1 between 62,437,899-62,438,699 bp) which belongs to a family protein that regulates different plant organs shape, including cotyledons, leaves, and fruits (Snouffer *et al*., 2020), and the *AP2-like* ethylene-responsive transcription factor *AINTEGUMENTA* or *ANT* gene (Solyc04g077490.3.1 between 60,418,478-60,421,941 bp), which plays a role as an auxin regulator in shoot and flower meristem maintenance, organ size and polarity, flower initiation, ovule development, floral organ identity, cell proliferation (Horstman *et al*., 2014). No high-effect variants were predicted by SnpEff in the coding sequence of these genes contrasting for the different founders’ phenotypes.

Haplotype results revealed a significant difference between SLC and SP founders according to pairwise *t*-test (Figure 4D). Although the haplotypes density plot also did not show a bimodal distribution for SLC and SP founders, it showed a higher density for SP haplotypes in lines exhibiting lack of lobing/serration or moderate lobing values, and a slightly higher density for SLC haplotypes in the very serrated leaf values.

#### Leaf complexity

The Manhattan plot for leaf complexity revealed one significant peak on chromosome 4 (Figure S5A, Table 3). For the GLM model, two SNPs were above the Bonferroni threshold (LOD > 5.11) between 62.49 and 62.73 Mb (Figure S5B). For the MLM and BLINK model, a single SNP was above the Bonferroni threshold (LOD = 5.38 and 8.93, respectively) at 62.49 Mb position. This association peak accounted for 4.12% of the total phenotypic variance of the leaf complexity trait.

Two genes involved in the leaf complexity were detected within the candidate genomic region on chromosome 4 identified in the GWAS (Table 3). In order of proximity to the candidate region we found the *KNOTTED1* gene (Solyc 04g077210.3.1 between 60,124,504-60,131,770 bp) which is expressed during leaf development and affects leaf morphology altering leaf complexity (Shani *et al.,* 2009), and the *entire* or *INDOLE-3-ACETIC ACID9 IAA9* gene (Solyc04g076850.3.1 between 59,750,087-59,755,552 bp), which controls leaf morphology from compound to simple leaves (Zhang *et al.,* 2007). No high-effect variants were predicted by SnpEff in the coding sequence of these genes for the founders with contrasting phenotypes.

Haplotype results revealed a significant difference between SLC and SP founders according to the pairwise *t*-test (Figure 4E). Although the haplotypes density plot did not show a bimodal distribution for SLC and SP founders, it showed a higher density for SP haplotypes in pinnate leaves, and a slightly higher density for SLC haplotypes in the bipinnate leaves.

### Earliness

The Manhattan plot for the number of leaves below the first inflorescence revealed one significant peak on chromosome 11 (Figure S6A, Table 3). For the GLM model, four SNPs were above the FDR and Bonferroni thresholds (LOD > 4.75 and 5.11, respectively) between 2.05 and 2.80 Mb. For the MLM model, only two SNPs were above the FDR and Bonferroni thresholds between a reduced region of 2.17 and 2.80 Mb (Figure S6B). For the BLINK model, a single SNP was above the FDR and Bonferroni thresholds (LOD = 24.22) at 2.80 Mb position. The association peak explained 5.52% of the total phenotypic variance of the number of leaves below the first inflorescence trait.

Different genes implicated in the flowering pathway were identified in the candidate genomic region on chromosome 11 proposed in the GWAS for the number of leaves below the first inflorescence (Table 3). In order of proximity to the candidate region we found two *FLOWERING LOCUS T* (*FT*) genes (*FT1* Solyc11g008640.1.1 between 2,854,837-2,857,237 bp and *FT2* Solyc11g008650.1.1 between 2,866,945-2,867,166 bp), which have been described as mediating the onset of flowering and the floral transition in all angiosperms (Pin and Nilsson, 2012), the *SELF-PRUNING INTERACTING PROTEIN 1* or *SP1* gene (Solyc11g007880.1.1 between 2,135,303-2,135,602 bp), which is involved in a conserved signalling system that regulates flowering (Pnueli *et al.,* 2001), and the *JOINTLESS* or *J* gene (Solyc11g010570.2.1 between 3,671,232-3,676,350 bp), which plays a role in flowering promotion (Szymkowiak and Irish, 2006). The FT1 and FT2 proteins have respectively a 71.68% (124/173) and 87.69% (57/65) identity with the well-known *SINGLE-FLOWER TRUSS* (*SFT*, Solyc03g063100.2.1) gene product according to BLASTp alignment. While *FT1* is recognized as a paralogue of the *SFT* gene in EnsemblPlants, *FT2* seems to be a truncated pseudogene. Nevertheless, no clear variants were predicted by SnpEff in the coding sequence of these genes contrasting for the different founders’ phenotypes.

Haplotype results did not differentiate between SLC and SP founders (Figure 4F). Pairwise *t*-test only revealed a significant difference between SLC1, SLC3, and SLC4 from SLC2, SP1, and SP2 founders, with SP3 and SP4 in intermediate positions. The haplotype density plot also did not show a bimodal distribution for SLC and SP founders. However, it showed a trend for lower number leaves below the first inflorescences for the SP haplotypes, while SLC haplotypes were distributed along a wide range of number of leaves below the first inflorescence.

## Discussion

We present a novel inter-specific ToMAGIC population of 354 lines constructed by combining the genomes of SLC and SP founders. SLC accessions are phylogenetically positioned between SP and cultivated tomato (Blanca *et al.,* 2015, 2022). Therefore, founders were selected to exploit the wide diversity found in the tomato closest relatives taking advantage of their interbreeding compatibility (Peralta *et al.,* 2008). Previous resequencing of the selected founders allowed to significantly enhance recombination detection, haplotype prediction, and causal variants identification within the MAGIC population (Gramazio *et al.,* 2020a).

The MAGIC population was generated through a systematic “funnel” approach (Arrones *et al*., 2020) involving multiple rounds of inter-cross of the eight selected founders and five generations of selfing, totalling ten generations. The three inter-crossing generations from the two double hybrids and the blind SSD process ensured high levels of recombination, maintaining a high genetic and morphological diversity. The ToMAGIC lines were genotyped by using a newly developed 12k probes tomato panel, based on SPET, which is a robust technology based on target SNPs, but also capable of discovering novel SNPs (Barchi *et al.,* 2019). Although SPET has been mostly used in the biomedical field, it has demonstrated its potential as a high-throughput and high-efficiency genotyping platform in *Solanum* species (Gramazio *et al.,* 2020b; Mangino *et al.,* 2022). In this study, more than 4 million SNPs were generated with the 12k probes tomato SPET panel. After stringent filtering, 6,488 were retained as markers, while in the previous tomato MAGIC population developed by Pascual *et al*. (2015), 1,486 markers obtained by a custom-made genotyping platform (Fluidigm 96.96 Dynamic Arrays, San Francisco, CA) were used for population analyses. The genotypic data revealed the absence of genetic structure, which is one of the advantages of MAGIC populations (Arrones *et al.,* 2020), and a balanced representation of the founder genomes. The average contribution of each founder to the overall population was around 12.50%, which is the expected value for a population developed from eight founders.

We have demonstrated the power of our ToMAGIC population for the fine mapping of traits of interest in tomato breeding. Specifically, GWAS analysis detected strong associations for all the traits evaluated using three different models (GLM, MLM, and BLINK), supporting the robustness of the associations detected (Price *et al.,* 2006; Yu *et al.,* 2006; Huang *et al.,* 2019).

The implementation of SLC and SP accessions as founders have introduced a wide genetic and phenotypic diversity in the ToMAGIC population (Blanca *et al.,* 2015; Gramazio *et al*., 2020a). Our proof-of-concept, focusing on a subset of traits from different plant parts has revealed a large phenotypic diversity in the ToMAGIC population, including transgressive lines to some of the founders for all traits except leaf morphology. Within the phenotypic diversity of the final population, wild alleles showed a dominant effect over domesticated alleles in most traits. For instance, ToMAGIC lines tend to produce small fruits and simpler leaves, more similar to SP than to cultivated tomato. This prevalent dominance of wild alleles has been previously observed during the development of other inter-specific populations (Semel *et al.,* 2006).

Large tomato fruit size is a typical domestication trait, controlled by at least five different genes (Pereira *et al.,* 2021). It is tempting to speculate that, similar to the non-shattering spike trait in cereals (Lin *et al.,* 2012), it negatively affects plant fitness in the wild, by reducing seed dispersal by small vertebrates. Drawing on this parallel, the most likely scenario is that recessive alleles for large fruit size in tomato and non-shattering spike in cereals were both pre-existing in wild/weedy populations, and that they were not completely counterselected due to their recessive nature. Under this scenario, human selection for higher harvestable biomass probably acted on the rare homozygous plants that appeared in these wild populations. Consistent with this hypothesis, the nonfunctional (domesticated) allele of the rice shattering gene *sh4* is found, at low frequency, in the wild ancestor *O. rufipogon* (Lin *et al.,* 2007).

Almost all wild tomato species produce bilocular small fruits, and therefore, locule number and fruit weight played a crucial role in the increase in fruit size during domestication (Alpert *et al*., 1995; Lippman and Tanksley, 2001; Barrero *et al.,* 2006). On one hand, as a result of the GWAS analysis for locule number, an associated genomic region was identified that colocalized with the *WUSCHEL* gene. Mutations on this gene have been necessary to increase locule number during domestication (Muños *et al.,* 2011). However, previous sequence analysis on this gene revealed that the diversity of this locus was drastically reduced in the cultivated species (Muños *et al.,* 2011; van der Knaap *et al.,* 2014). Only two SNPs have been identified in this gene responsible for the large-fruited phenotype, which are the same two SNPs that we have found in our population. On the other hand, the GWAS analysis for fruit weight revealed an associated genomic region on chromosome 2 between 50.51 and 50.55 Mb in the region where the *FW2.2* gene is located (Frary *et al.,* 2000). Similarly, but not as precisely as in our ToMAGIC populations, in the tomato MAGIC developed by Pascual *et al*. (2015) a peak with the highest LOD value between 46.35 and 47.49 Mb was also identified. The *FW2.2* gene is responsible for up to 30% of the fruit weight variation between large domesticated tomatoes and the small-fruited wild relatives (Nesbitt and Tanksley, 2001). All modern tomatoes contain the large-fruited allele for *FW2.2* (Blanca *et al.,* 2015; Beauchet *et al.,* 2021), which was also identified in the two large-fruited SLC ToMAGIC founders. Molecular evolutionary studies suggested that this allele originated in wild tomatoes long before the process of domestication (Nesbitt and Tanksley, 2002). Indeed, fruit weight was strongly selected in SLC in the Andean region of Ecuador and Northern Peru prior to the domestication of tomato in Mesoamerica (Blanca *et al.,* 2015).

Anthocyanins are the main responsible for purple pigmentation in tomato leaf veins, leaf tissues, and stem (Barrett *et al.,* 2010; Jaakola, 2013). Plant anthocyanins are more commonly present in wild tomato species, where they have a main protective function against UV-visible light and other stressful conditions such as cold temperature, pathogens, or drought (Gould, 2004; Olsen *et al*., 2009; Zhang *et al*., 2014). The GWAS results identified an associated genomic region which colocalized with the previously described *SlMYB-ATV* gene. Overexpression of the coding protein acts as an inhibitor of anthocyanin production by silencing key regulators of the biosynthesis pathway (Cao *et al*., 2017; Colanero *et al*., 2018). The *atv* mutation was described as a 4 bp insertion in the second exon which led to a frameshift variant resulting in a premature stop codon with a strong impact in the polypeptide. This mutation was identified as the causal agent of anthocyanin production in the vegetative part of the plant (Colanero *et al*., 2018). Here, a novel mutation in the “purple” SP4 founder was found. Specifically, a 9 bp deletion leading to a disruptive inframe deletion which directly affects the transcription repressor MYB domain was identified. This demonstrates the significance of the ToMAGIC population as a reservoir of novel candidate genes and causative alleles. Interestingly, of the four SP founders, SP4 is the only one showing anthocyanin pigmentation as well as the one collected at the highest altitude (1,020 m) and lowest mean annual temperature (13°C), in agreement with the proposed role of anthocyanins as UV-sunscreens in cold temperatures (Martínez-Cuenca *et al.,* 2020).

Cultivated tomato leaf morphology has typical bipinnate compound leaves with moderately deep lobes, while there is a huge diversity of leaf morphology among wild tomato species (Zhang *et al.,* 2007; Kang *et al.,* 2010; Nakayuma *et al.,* 2023). Since leaf lobing/serration and leaf complexity traits are correlated, both traits have usually been studied together (Kang *et al.,* 2010). Actually, the GWAS results identified an associated genomic region on chromosome 4 around 62 Mb position for both traits, and candidate genes affecting both traits were identified within this genomic region. Although the *AP3/DEF* gene has mainly been related to petal and sepal development, other genes belonging to the same MADS box family are involved in tomato leaf development. Specifically, the *APETALA1/FRUITFULL* (*AP1/FUL*) MADS box genes are involved in the organogenic activity of the leaf margin and leaf complexity (Burko *et al.,* 2013). The *ANT* gene also belongs to a family of APETALA 2/ETHYLENE RESPONSE FACTOR (AP2/ERF) domain transcription factors which affects plant leaf shape and size by regulating cell proliferation (Horstman *et al.,* 2014). The *OVATE* gene was first identified in tomato as a key regulator of fruit shape (Wang *et al.,* 2016). However, expression of *OVATE* genes can also result in dwarf plants with shorter and thicker organs such as rounder leaves (Snouffer *et al.,* 2020). The tomato *KNOTTED1* promotes cytokinin biosynthesis which is directly related to cell proliferation (Nakayuma *et al.,* 2023), and different levels of cytokinins led to a broad spectrum in leaf complexity (Shani *et al.,* 2009; Shwartz *et al.,* 2016). This gene has a key role in the molecular mechanism behind leaf development and evolution and has been repeatedly exploited to generate natural variations in leaf shape (Ichihashi and Tsukaya, 2015). The *IAA9* gene is a transcriptional repressor in auxin signal transduction (Abe-Hara *et al.,* 2021). Tomato mutants for *IAA9* also showed altered leaf morphology with the compound leaf changing to a single leaf (Zhang *et al.,* 2007; Ueta *et al.,* 2017; Abe-Hara *et al.,* 2021). In this way, leaf development is mainly influenced by cell proliferation and different hormones as a result of the activity of a complex gene network (Nakayuma *et al.,* 2023). An accurate phenotyping of the ToMAGIC population for these traits has allowed to narrow down a genomic region that harbours a large number of genes related to leaf morphology. This genomic region could be further narrowed down by studying the segregation of the cross between two isolines to enable the identification of the responsible gene/s.

The existence of early-flowering alleles in wild species indicates the relevance of exploiting the genetic variation present in tomato wild relatives (Jiménez-Gómez *et al.,* 2007). Although the mechanisms controlling the transition from vegetative to reproductive growth are complex, several genes involved in flowering regulation are known (Meir *et al.,* 2021; Zhang *et al.,* 2023b). The number of leaves below the first inflorescence trait is a proxy for earliness in tomato (Honma *et al.,* 1963) and is easily scored and commonly assessed to evaluate the earliness in tomato (Jiménez-Gómez *et al.,* 2007; Nakano *et al.,* 2016; Silva *et al.,* 2019). The GWAS analysis for the number of leaves below the first inflorescence identified an association on chromosome 11, where several genes related to flowering time were found (two *FT* genes, *SP1*, and *J*). The most studied *FT* gene is the tomato ortholog *SINGLE-FLOWER TRUSS* (*SFT*) gene on chromosome 3, which encodes for florigen and induces flowering in day-neutral (Turck *et al.,* 2008; Meir *et al.,* 2021; Zhang *et al.,* 2023b). Here, we report the *FT1* gene on chromosome 11, a paralogue of the *SFT* gene which may also be involved in the flowering regulation. The *SP1* gene is a member of the *CETS* family of regulatory genes, together with *FT* genes, controlling flowering time (Pnueli *et al.,* 2001). However, they play an antagonistic role, since *SP1* delays flowering in tomato (Zhang *et al.,* 2023b). The *J* gene is involved in the same pathway as the *SFT* gene but with a small role in flowering promotion (Szymkowiak and Irish, 2006; Zhang *et al.,* 2023b). A better understanding of the mechanisms underlying the tomato flowering regulatory pathways will allow breeding to target more precise candidate genes for the induction of early flowering. Nevertheless, once again, the ToMAGIC population has led us to a genomic region directly involved in the transition to flowering, pointing to new candidate genes.

Overall, the genotyping results together with the large morphological variation observed in the new inter-specific SLC/SP tomato MAGIC population, as well as the appearance of transgressive phenotypes, indicate that recombination and variation were maximised in the final population. The ToMAGIC population has demonstrated a high potential for the fine mapping of traits of interest from different plant parts. Given the fact that the population contains representatives of the tomato ancestor (SP) and the primitive weedy forms (SLC) of tomato, it can also be a tool of great relevance for studying the genetic changes in the early stages of tomato domestication. It is also evident from our study that the derived ToMAGIC population or core collections developed from it can contribute to tomato genetics research and breeding programs. Recombinant lines with combinations of traits of interest present in different founders can also be of direct interest to breeders or even for selection of small-fruited new cultivars.

## Acknowledgements

Funding for this work has been received from the following funders: MCIN/AEI/10.13039/501100011033 (grant PID2020-118627RB-I00), MCIN/AEI /10.13039/501100011033 and European Union Europea NextGenerationEU/ PRTR (grant TED2021-129296B-I00), Conselleria d’Innovació, Universitats, Ciència i Societat Digital of the Generalitat Valenciana (grant CIPROM/2021/020), European Commission H2020 Research and Innovation Programme through the HARNESSTOM innovation action (grant agreement No. 101000716) and the Horizon Europe PRO-GRACE project (grant agreement No. 10194738). Andrea Arrones is grateful to Spanish Ministerio de Ciencia, Innovación y Universidades for a predoctoral (FPU18/01742) contract. Oussama Antar is grateful to Conselleria d’Innovació, Ciència i Societat Digital of the Generalitat Valenciana for a pre-doctoral grant within the Santiago Grisolía program (CIGRIS/2022/113). Leandro Pereira-Dias is grateful to Universitat Politècnica de Valencia and the Spanish Ministerio de Universidades for a post-doctoral grant under the Margarita Salas funded by the European Union NextGenerationEU/PRTR. Pietro Gramazio is grateful to Spanish Ministerio de Ciencia e Innovación for a post-doctoral grant (RYC2021-031999-I) funded by MCIN/AEI/10.13039/501100011033 and the European Union through NextGenerationEU/PRTR.

## Contributions

SV, PG, MJD, and JP conceived the idea and supervised the manuscript; AA OA, LP-D, AS, and MJD performed the field trials. GA and GG in collaboration with TECAN Genomics designed the 12k SPET panel. All authors analysed the results. AA and OA prepared a first draft of the manuscript and the rest of authors reviewed and edited the manuscript. All authors have read and agreed to the published version of the manuscript.

## Data availability statement

The datasets presented in this study can be found in online repositories. The names of the repository/repositories and accession number(s) can be found below: https://www.ncbi.nlm.nih.gov/, PRJNA616074.

## Supplementary information

**Figure S1.** (A) A representation of different phenotypes for the locule number, fruit weight, and plant anthocyanin traits, together with the known genes controlling these traits indicating the phenotypic-causing variants. (B) Correlation analysis among all the studied traits showing a slight positive correlation between the leaf morphology traits, corresponding to leaf lobing/serration and leaf complexity. On the right, a representation of the phenotypic scores for both traits.

**Figure S2.** Genome-wide association results for the fruit weight trait. (A) Manhattan plots comparing GLM, MLM and BLINK models. (B) On the top, a chromosome-wise Manhattan plot with the top significant markers. Bonferroni threshold is represented with red dashed line. On the bottom, heat map of pairwise linkage disequilibrium (LD).

**Figure S3.** Genome-wide association results for the plant anthocyanin trait. (A) Manhattan plots comparing GLM, MLM and BLINK models. (B) On the top, a chromosome-wise Manhattan plot with the top significant markers. Bonferroni and FDR thresholds are represented with red dashed and continuous lines, respectively. On the bottom, heat map of pairwise linkage disequilibrium (LD).

**Figure S4.** Genome-wide association results for the leaf lobing/serration trait. (A) Manhattan plots comparing GLM, MLM and BLINK models. (B) On the top, a chromosome-wise Manhattan plot with the top significant markers. Bonferroni and FDR thresholds are represented with red dashed and continuous lines, respectively. On the bottom, heat map of pairwise linkage disequilibrium (LD).

**Figure S5.** Genome-wide association results for the leaf complexity trait. (A) Manhattan plots comparing GLM, MLM and BLINK models. (B) On the top, a chromosome-wise Manhattan plot with the top significant markers. Bonferroni threshold is represented with red dashed line. On the bottom, heat map of pairwise linkage disequilibrium (LD).

**Figure S6.** Genome-wide association results for the number of leaves below the first inflorescence trait. (A) Manhattan plots comparing GLM, MLM and BLINK models. (B) On the top, a chromosome-wise Manhattan plot with the top significant markers. Bonferroni and FDR thresholds are represented with red dashed and continuous lines, respectively. On the bottom, heat map of pairwise linkage disequilibrium (LD).

