## Supplementary figures and images for "A novel tomato inter-specific (*Solanum lycopersicum* var. *cerasiforme* and *S. pimpinellifolium*) MAGIC population facilitates trait association and candidate gene discovery in untapped exotic germplasm"

### Figure S1

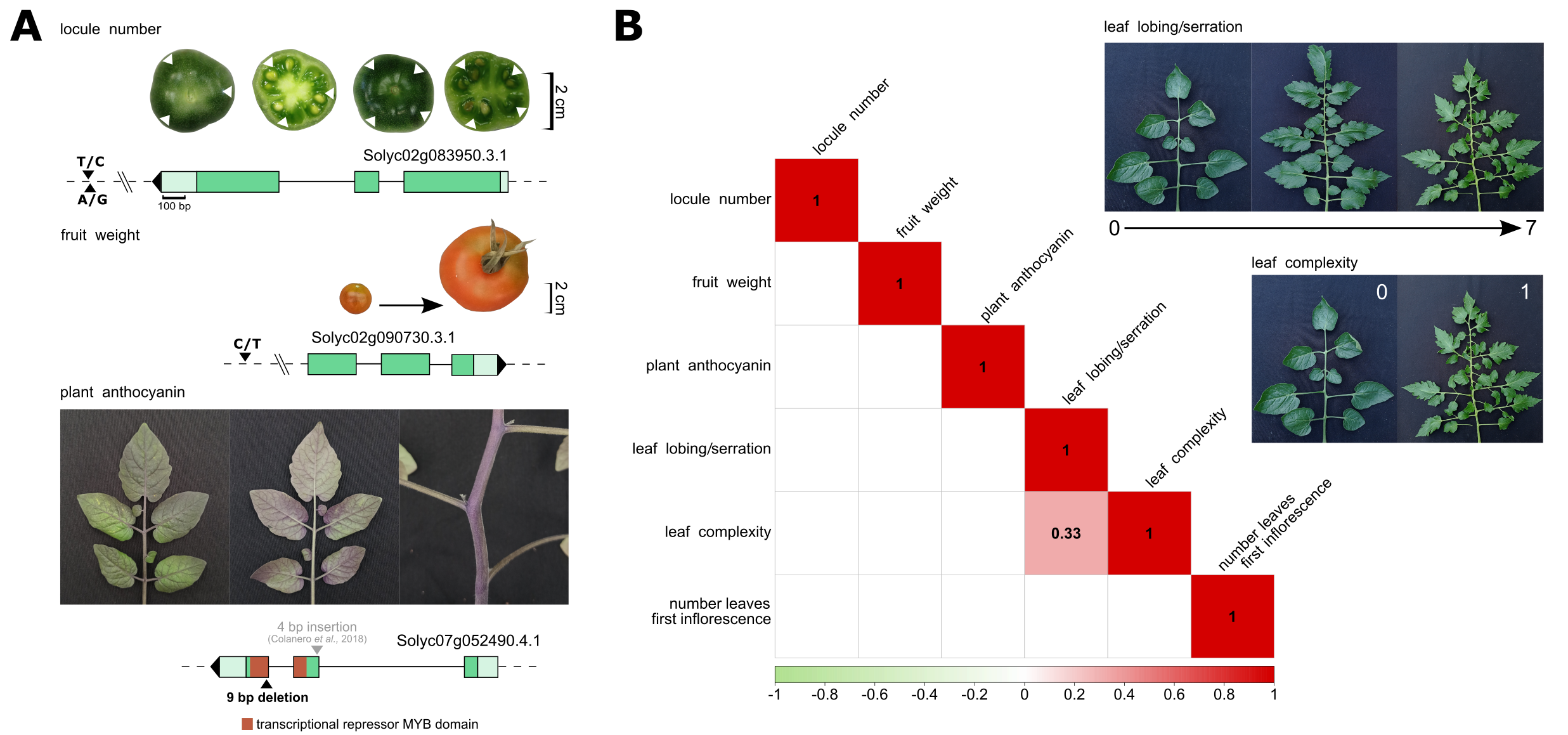

### Figure S3

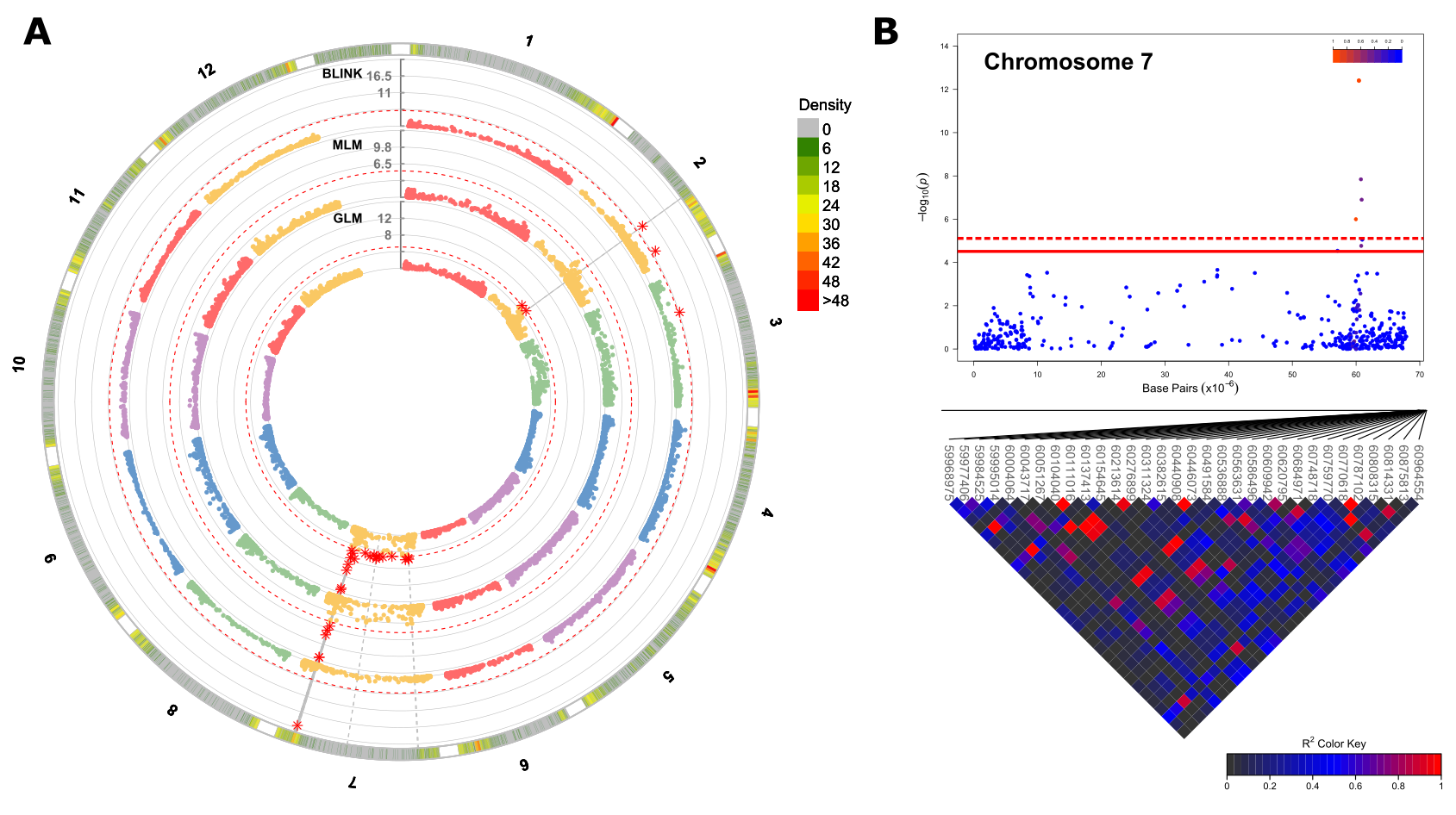
